# *Borrelia burgdorferi* DnaA and the nucleoid-associated protein EbfC coordinate expression of the *dnaX-ebfC* operon

**DOI:** 10.1101/2022.10.17.512582

**Authors:** Andrew C. Krusenstjerna, William K. Arnold, Timothy C. Saylor, Jamila S. Tucker, Brian Stevenson

## Abstract

*Borrelia burgdorferi*, the spirochete agent of Lyme disease, has evolved within a consistent infectious cycle between tick and vertebrate hosts. The transmission of the pathogen from tick to vertebrate is characterized by rapid replication and a change in the outer surface protein profile. EbfC, a highly conserved nucleoid-associated protein, binds throughout the borrelial genome affecting expression of many genes, including the Erp outer surface proteins. In *B. burgdorferi*, like many other bacterial species, *ebfC* is co-transcribed with *dnaX*, an essential component of the DNA polymerase III holoenzyme, which facilitates chromosomal replication. The expression of the *dnaX-ebfC* operon is tied to the spirochete’s replication rate, but the underlying mechanism for this connection was unknown. In this work, we provide evidence that the expression of *dnaX-ebfC* is controlled by direct interactions of DnaA, the chromosomal replication initiator, and EbfC at the unusually long *dnaX-ebfC* 5’ UTR region. Both proteins bind to the 5’ UTR DNA, with EbfC also binding to the RNA. The DNA binding of DnaA to this region was similarly impacted by ATP/ADP. *In vitro* studies characterized DnaA as an activator of *dnaX-ebfC* and EbfC as an anti-activator. We further found evidence that DnaA may regulate other genes essential for replication.

**IMPORTANCE:** The dual-life cycle of *Borrelia burgdorferi*, the causative agent of Lyme disease, is characterized by periods of rapid and slowed replication. The expression patterns of many of the spirochete’s virulence factors are impacted by these changes in replication rates. The connection between replication and virulence can be understood at the *dnaX-ebfC* operon. DnaX is a component of the DNA polymerase III holoenzyme that facilitates replication. EbfC is a nucleoid-associated protein that regulates the infection-associated outer surface Erp proteins, as well as other transcripts. The expression of *dnaX-ebfC* is tied to replication rate, which we demonstrate is mediated by DnaA, the master chromosomal initiator protein and transcription factor, and EbfC.

## INTRODUCTION

Replication of the chromosome is a crucial yet costly process. Across all domains of life, chromosomal replication has evolved to require an initiator protein and a replicase (1, 2). For bacteria, the DNA-binding protein DnaA and the DNA polymerase III holoenzyme (DNA Pol III HE) fulfill these roles, respectively. DnaA associates at the chromosomal origin of replication (*oriC*) to high-affinity DnaA-boxes and, through cooperative oligomerization to low-affinity DnaA-boxes, melts the DNA to allow for loading of the helicase and the DNA Pol III HE (3–5). As expected, bacteria have developed distinct pathways to guarantee that this process is initiated appropriately for its environmental conditions.

The dual-host enzootic lifecycle of *Borrelia burgdorferi* is intimately tied to replication. This connection is best understood at the tick-vertebrate transmission interface. In the mid-gut of an unfed tick, spirochetes do not replicate, but this changes when the vector takes a blood meal (6–8). Increased nutrient availability permits the spirochetes to replicate their genomes and alter their proteomes to permit the transmission from tick to vertebrate. For *B. burgdorferi*, many of these host-specific phenotypic switches are regulated by a limited repertoire of nucleic acid-binding proteins (9). A notable example is the nucleoid-associated protein EbfC, which directly regulates the production of vertebrate-stage Erp outer surface proteins and numerous other transcripts (10, 11).

EbfC is highly conserved across eubacteria and is typically the second gene in a bicistronic operon with *dnaX*, an essential component of the DNA Pol III HE (12–14). In the model organism *Escherichia coli*, *dnaX* codes for two subunits, the full-length Tau (τ) and shortened Gamma (γ), by translational frameshift (15–17). All bacterial replicases are composed of three domains essential to their function: a core polymerase, a sliding processivity clamp, and a clamp loading complex. Tau is the central organizer of these tripartite bacterial replicases, connecting the core polymerases on the leading and lagging strands with the processivity clamp loading complex.

For many bacterial species, DnaA functions as a transcription factor coordinating the expression of genes for various cellular pathways (18, 19). In *B. burgdorferi*, *dnaX-ebfC* transcript levels are tied to bacterial replication rates (20, 21), which we hypothesized could be mediated by DnaA. We found that both DnaA and EbfC contribute to the expression of the *dnaX-ebfC* operon. Additional evidence suggests that DnaA might also be involved with the production of other components of DNA Pol III HE and DNA gyrase.

## RESULTS

### DnaA binds to *dnaX-ebfC* 5’ UTR DNA

Using promoter-reporter fusion constructs, we previously mapped the promoter for the *dnaX-ebfC* operon to be between −247 and −166 bp upstream of the *dnaX* start codon (20). This region is just within the end of the gene for the non-coding signal recognition particle RNA (*srp*; **Fig. 1A**). *In silico* bacterial promoter analysis (BPROM, Softberry; (22)) of this segment of DNA predicted a single σ^70^ promoter, indicating the *B. burgdorferi dnaX-ebfC* operon has a 5’ UTR of about 179 nucleotides (**Fig. 1B**). For context, the median 5’ UTR in *B. burgdorferi* is ∼36 nucleotides in length (23). This significant deviation, in conjunction with the importance of this operon’s products to borrelial biology, prompted us to hypothesize that the 5’ UTR is critical to *dnaX-ebfC* regulation.

**Figure 1.**
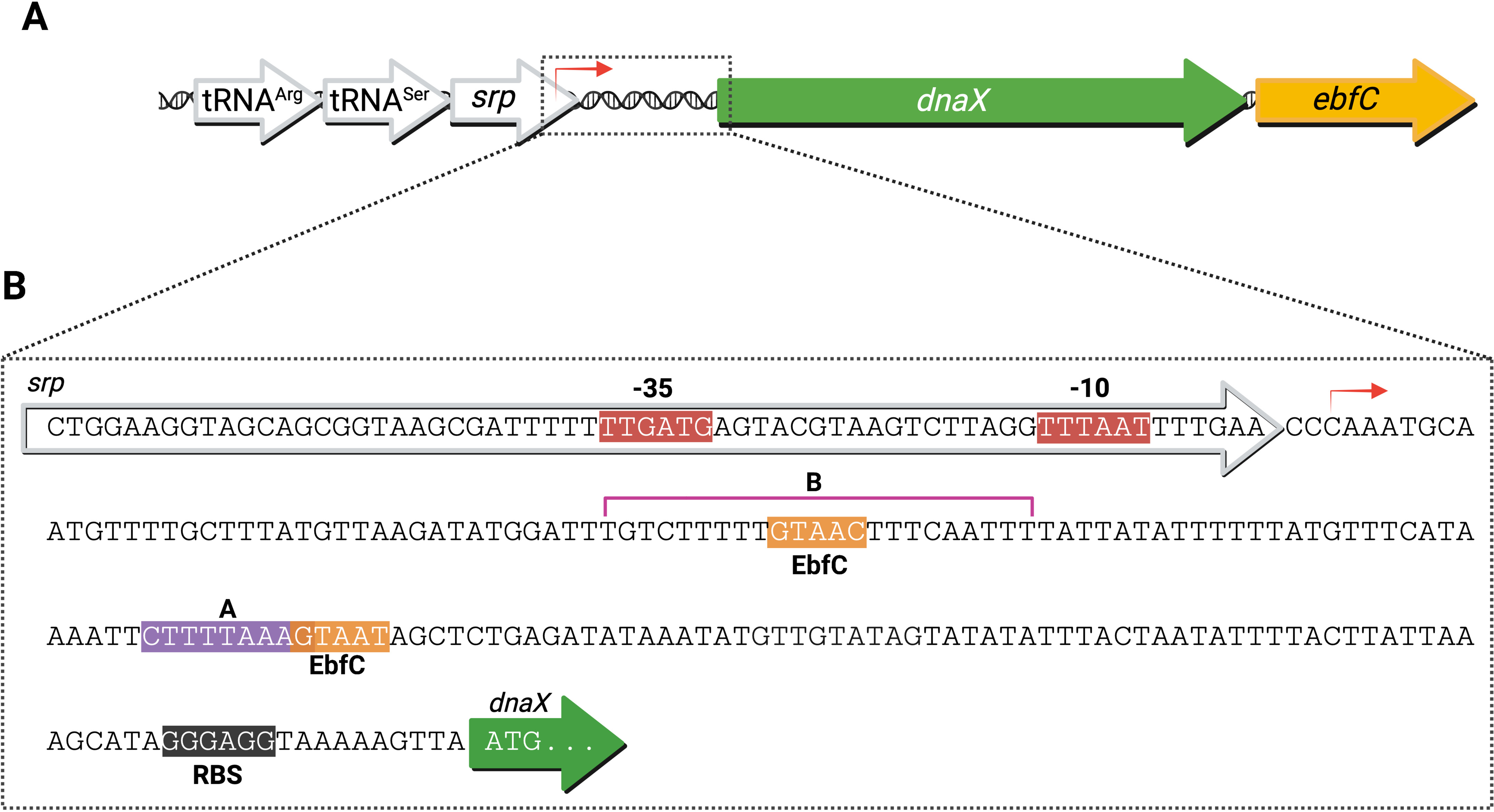
The *dnaX-ebfC* operon and its 5’ UTR. (A) The *B. burgdorferi dnaX-ebfC* locus is located downstream of genes for two tRNAs and the RNA component of the signal recognition particle (SRP). (B) The predicted promoter (Softberry, BPROM) of the *dnaX-ebfC* operon is just within the end of the *srp* gene. The 179 nt 5’ UTR contains a hypothesized DnaA-box (A), and two EbfC binding motifs. The region labeled B is the sequence DnaA was also found to bind (*see below*).

As DnaX and EbfC are evidently important for *B. burgdorferi* during periods of rapid replication, we first pursued whether DnaA, which is both the master replication initiator and a transcription factor in other species, can interact with *dnaX-ebfC*. Sequence level examination of the 5’ UTR DNA of *dnaX-ebfC* revealed a potential DnaA-box (**Fig. 1B**) (24). To test whether DnaA binds the *dnaX-ebfC* 5’ UTR DNA, we performed electrophoretic mobility shift assays (EMSAs) using a recombinant DnaA with a glutathione S-transferase (GST) solubility tag fused to the N-terminus (GST-DnaA).

In other bacterial species, the DNA binding activity of DnaA is affected by whether it has ATP or ADP-bound (25–27). As our recombinant GST-DnaA was purified under denaturing conditions and refolded without ATP/ADP, we surmised that the protein adopted an apo-form before use. Thus, our EMSA DNA-binding reactions with GST-DnaA were supplemented with saturating concentrations of either ATP or ADP (**Fig. 2A-B**). Under these conditions, we observed concentration-dependent binding of GST-DnaA to the 186 bp *dnaX-ebfC* 5’ UTR DNA probe. Control EMSAs with purified GST confirmed that this binding was not due to the solubility tag (**Fig. 2C**). Quantitative analysis of densitometric data from triplicate EMSAs with GST-DnaA gave overlapping apparent dissociations constants (K_D_^app^) of 1141 nM (95% CI: 703.5-1880 nM) and 737.7 nM (95% CI: 334.8-1639 nM) for ATP and ADP reactions, respectively (**Fig. 2D-F**). These differences were not statistically significant (*p* = 0.9835, unpaired student’s t-test). We concluded that DnaA binds the *dnaX-ebfC* 5’ UTR DNA and is not significantly impacted by added ATP/ADP.

**Figure 2.**
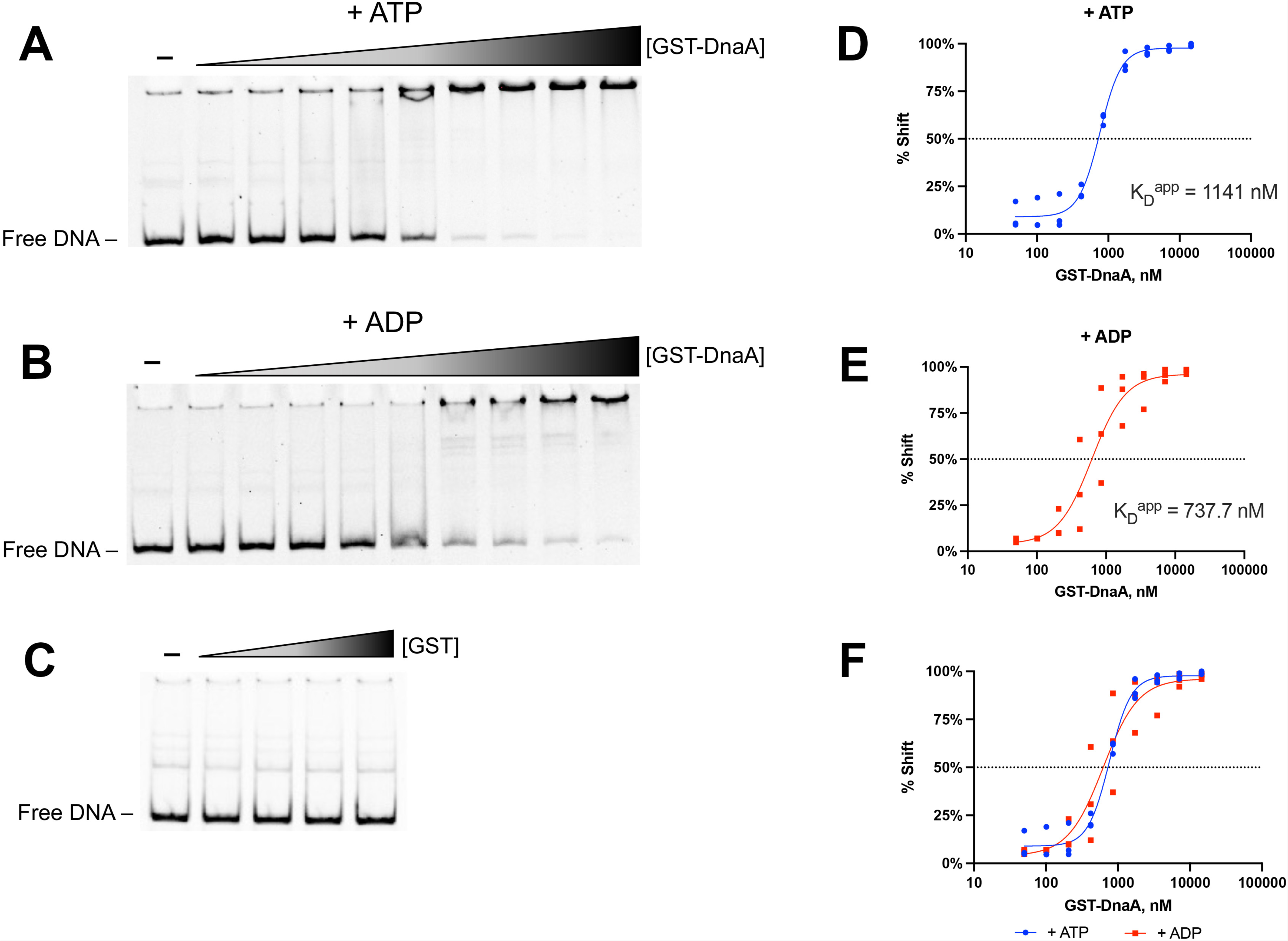
Interactions of DnaA at the *dnaX-ebfC* 5’ UTR DNA and assessment of effects of ATP and ADP. Electrophoretic mobility shift assays (EMSAs) were conducted with a fluorescent 186-mer DNA probe (100 nM) that consists of the *dnaX-ebfC* 5’ UTR. Representative EMSAs with 0.050, 0.102, 0.206, 0.419, 0.851, 1.728, 3.509, 7.126 and 14.5 µM of recombinant GST-DnaA with either 1 mM (A) ATP or (B) ADP. The first lanes contain the DNA probe only. (C) EMSA with 0.625, 1.25, 2.5, and 5 µM recombinant GST demonstrates that the observed GST-DnaA binding is not due to the fusion partner. (D-F) Apparent dissociation constant (K_D_^app^) measurements of GST-DnaA-probe interactions with ATP or ADP. Plots show the K_D_^app^ calculated using the densitometric data from triplicate EMSAs. The average K_D_^app^ for ATP reactions is 1141 nM and 737.7 nM for ADP. The difference in GST-DnaA-probe binding between the ATP and ADP reactions was not statistically significant (p = 0.9835, unpaired student’s t-test).

To narrow down where DnaA is binding in the *dnaX-ebfC* 5’ UTR DNA, a competition EMSA with unlabeled competitor DNAs was conducted. The DNA competitors were seven overlapping unlabeled 40-mer sequences from the labeled probe (**Fig. 3A**). The addition of 10-times molar excess of competitors 2 and 3 (C2 & C3), relative to the probe resulted in an appreciable increase in free DNA (**Fig. 3B**, lanes 6 and 7, respectively), indicating preferential binding of the protein to these regions. Competitor 3 contains the hypothesized DnaA-box, but competitor 2 (C2) does not contain a sequence resembling any previously characterized DnaA-box. The inability of the two DNAs that overlap competitor 2 (C5 and C6) to diminish the shifted complex indicates that DnaA bound to a region we term region B (**Fig. 3A**). As a positive control, a specific DNA competitor derived from the pCR2.1 TA clone containing the *dnaX-ebfC* 5’ UTR was used. The addition of this DNA resulted in the abrogation of the protein-DNA probe complex (**Fig. 3B**, PC). A negative control, the non-specific DNA derived from the empty pCR2.1 vector, did not affect the protein-probe complex (**Fig. 3B**, NC). These results demonstrate that DnaA binds specifically to regions A and B within the *dnaX-ebfC* 5’ UTR DNA.

**Figure 3.**
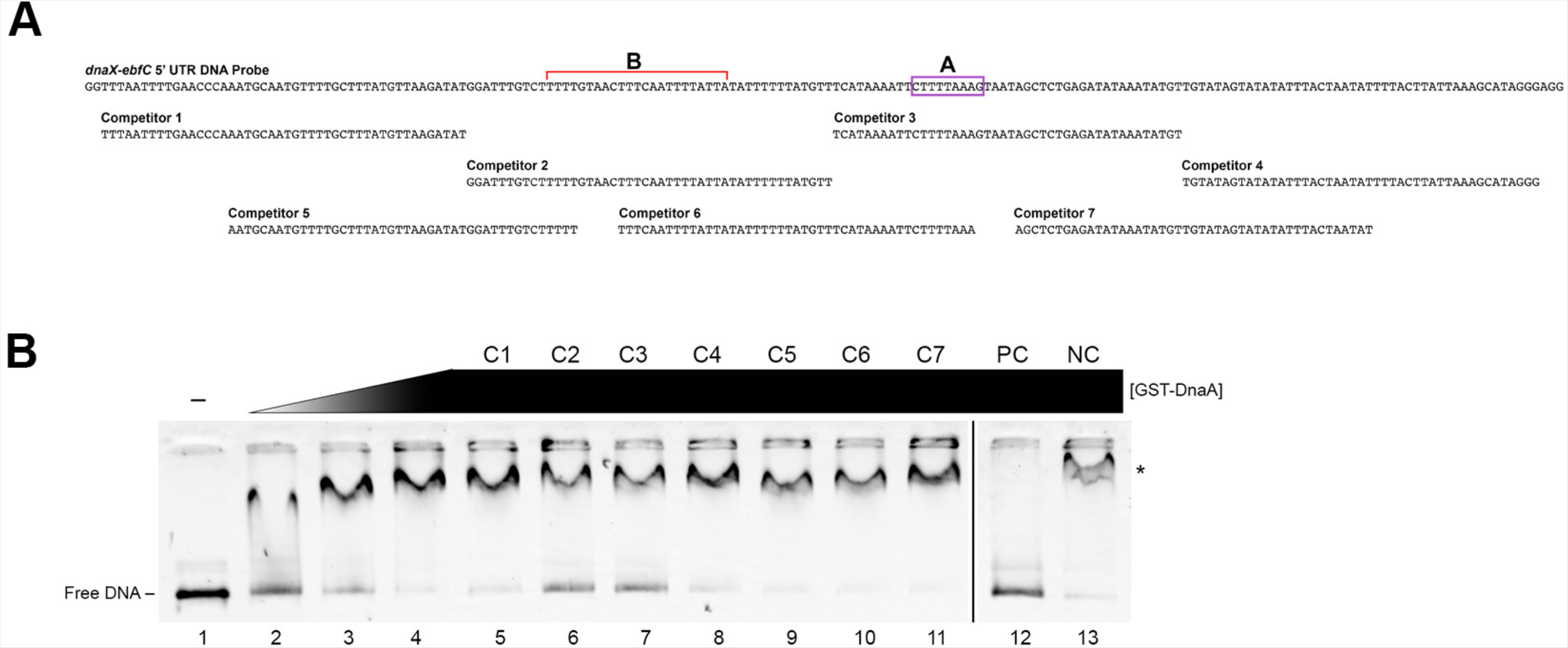
Identification of DnaA binding sites within the *dnaX-ebfC* 5’ UTR DNA. (A) Schematic of the 186-mer *dnaX-ebfC* 5’ UTR DNA probe and the seven overlapping 40-mer DNA competitors. Hypothesized DnaA-box is labeled A. The region where DnaA was also found to bind is labeled B. (B) Horizontal EMSA with a polyacrylamide TBE gel using 100 nM DNA probe, GST-DnaA, ATP, and different DNA competitors. Lane 1: probe only. Lanes 2-4: probe with 1.125, 2.25, and 4.5 µM GST-DnaA. Lanes 5-11: probe with 4.5 µM GST-DnaA and 10-molar excess of competitor DNAs (C1-C7). Lane 12: probe with 4.5 µM GST-DnaA and 10-molar excess of DNA amplified from pCR2.1 containing the *dnaX-ebfC* 5’UTR (Positive control, PC). Lane 13: probe with 4.5 µM GST-DnaA and 10-molar excess of DNA amplified from the empty pCR2.1 (Negative control, NC). The asterisk marks the probe-protein complex. The black line indicates the removal of extraneous lanes from the same gel.

### EbfC binds to the *dnaX-ebfC* 5’ UTR DNA and RNA

In addition to the DnaA-boxes identified upstream of the *dnaX* ORF, we also noted two EbfC binding motifs (**Fig. 1A**). EbfC binds with highest affinity to the broken palindromic sequence GTnAC and a lesser affinity to the partial sequence GTnAT (28). EMSAs with recombinant EbfC, nonspecific poly dI-dC competitor, and the 186-mer DNA probe showed specific binding of the protein to this region (**Fig. 4A**). Further EMSAs with 30-mer DNA probes containing either the consensus or partial EbfC motif verified that the EbfC bound each of those sequences (**Fig. 4B-C**). The binding to the complete motif probe was appreciably more prominent than the partial motif probe, consistent with previous characterizations. These results demonstrate that EbfC binds two sites within the *dnaX-ebfC* 5’ UTR DNA.

**Figure 4.**
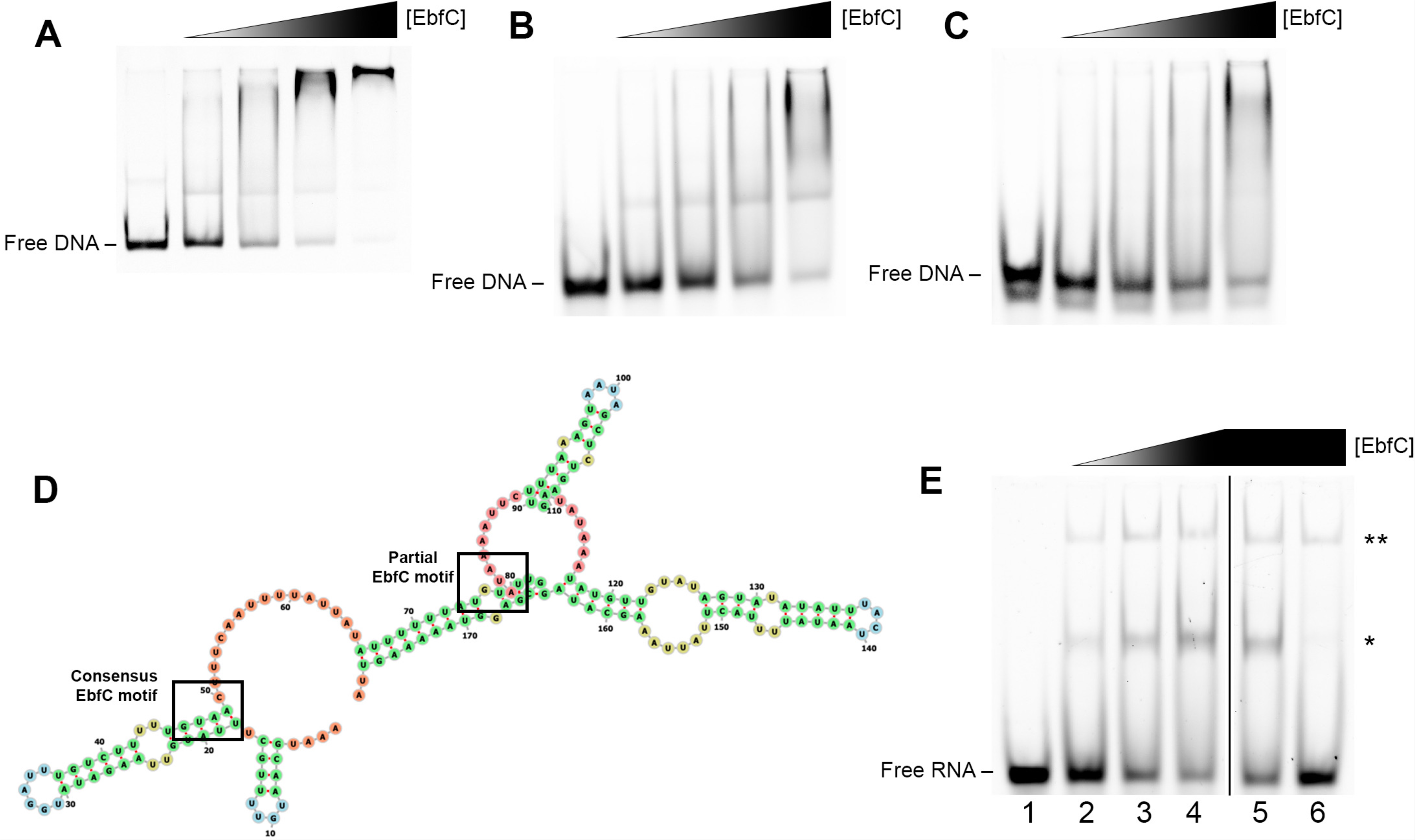
Interactions of EbfC with the *dnaX-ebfC* 5’ UTR DNA and RNA. (A-C) EMSAs with 100 nM labeled DNA probes, 2.5 ng/µl poly dI-dC, and either 3.75, 7.5, 15, and 30 µM recombinant EbfC. Lane 1 for each panel is probe only. (A) 186-mer DNA probe of the *dnaX-ebfC* 5’ UTR. (B) 30-mer DNA probe from the *dnaX-ebfC* 5’ UTR with the consensus EbfC motif. (C) 30-mer DNA probe from the *dnaX-ebfC* 5’ UTR with the partial EbfC motif. (D) Predicted RNA fold structure of the 179 nt 5’ UTR of *dnaX-ebfC* (RNAfold, http://rna.tbi.univie.ac.at/cgi-bin/RNAWebSuite/RNAfold.cgi). The consensus EbfC binding motif is predicted to be part of a stem-loop, and the partial EbfC binding motif is within a knot. (E) EMSA with 20 nM EbfC a 30-mer Alexa 647 RNA probe containing the consensus EbfC binding motif from the *dnaX-ebfC* 5’ UTR and recombinant EbfC. Lane 1: probe only. Lanes 2-4: probe with 5, 10 and 15 µm EbfC. Lane 5: probe with 1 µM competitor *flaB* RNA, and 15 µM EbfC. Lane 6: probe with 1 µM competitor *dnaX-ebfC* RNA. The lower-order complex is indicated by one asterisk (*), and the higher-order complex by two asterisks (**). All lanes are from the same gel, with the black line indicating the removal of extraneous lanes.

During our studies, colleagues shared that EbfC could be pulled down with total RNA from extracts of *B. burgdorferi* (29). We hypothesized that EbfC might also bind the extended *dnaX-ebfC* 5’ UTR RNA (**Fig. 4D**). To examine this, we performed EMSAs with a 30-mer RNA probe containing the consensus EbfC motif from the *dnaX-ebfC* 5’ UTR. Reactions with the RNA probe and recombinant EbfC resulted in the formation of two RNA-protein complexes (**Fig. 4E**, lanes 2-4). The addition of 50-times molar excess of RNA from the constitutively expressed *flaB* did not compete away the complexes (**Fig. 4E**, lane 5). At the same molar excess, the unlabeled *dnaX-ebfC* RNA successfully eliminated the smaller complex (**Fig. 4E**, lane 6). These results demonstrated that EbfC can bind untranslated RNA 5’ of the *dnaX-ebfC* open reading frames.

### Assessment of simultaneous binding of DnaA and EbfC to the *dnaX-ebfC* 5’ UTR DNA

Having established that DnaA and EbfC bind the 5’ UTR DNA of *dnaX-ebfC*, we sought to determine how the proteins would interact in this region. To examine this, EMSA reactions with constant EbfC and increasing GST-DnaA were prepared (**Fig. 5A)**. The constant EbfC concentration allowed for roughly equivalent levels of bound and unbound DNA (**Fig. 5A**, lane A). With increasing concentrations of GST-DnaA (**Fig. 5A**, lanes B-G), the DNA-protein complex migrated more slowly in the gel. Densitometry showed an initial increase of free DNAs at low levels of added GST-DnaA (**Fig. 5B**, lanes B-D), followed by a decrease at the highest GST-DnaA concentrations (**Fig. 5B**, lanes E-G). This pattern was not appreciably affected by ATP/ADP. These data suggest that the two proteins may bind cooperatively to the *dnaX-ebfC* 5’ UTR DNA. The precise stoichiometry and kinetics of the DNA interactions are undoubtedly complex, given the proximity of the binding sites for each protein.

**Figure 5.**
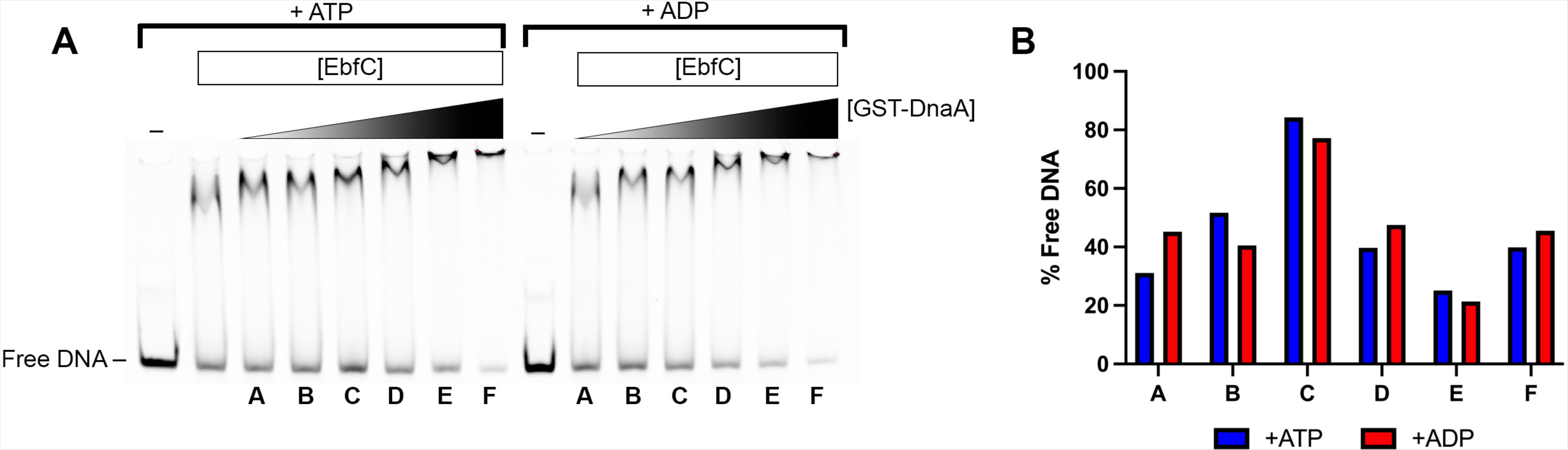
Interaction of DnaA and EbfC at the *dnaX-ebfC* 5’ UTR DNA. (A) EMSA using the *dnaX-ebfC* probe with constant EbfC concentration of 10 µM and 0.078, 0.156, 0.312, 0.625, 1.25, and 2.50 µM of GST-DnaA. Reaction A is EbfC only with the probe. Reactions B-G have an increasing concentration of GST-DnaA and constant EbfC concentration. The first eight lanes include 1 mM ATP, and the last seven include 1 mM ADP. (B) Graph of the percent free DNA for each reaction as determined by densitometry. For both ATP and ADP conditions, free DNA increased (reaction D) at lower concentrations of added GST-DnaA. At the highest GST-DnaA concentrations, free DNA decreased.

### DnaA enhances the expression of *dnaX-ebfC*, while EbfC inhibits this effect

Having established the interactions of DnaA and EbfC at the *dnaX-ebfC* DNA/RNA, the next logical step was to determine whether the proteins affect the expression of *dnaX-ebfC*. As both proteins are essential for the survival of *B. burgdorferi* (28), analyses with deletion mutant bacteria could not be done. Instead, cell-free *in vitro* transcription/translation assays with *E. coli* S30 extracts were performed. This method has been a useful tool for quantifying DNA-binding proteins’ effects on borrelial gene and protein expression and has the added benefit of lacking *B. burgdorferi* DnaA or EbfC (10, 11, 30). The template for these reactions was a linear DNA that contains the 247 bp 5’ of *dnaX,* including the transcriptional promoter, fused to *gfp* (20). The addition of GST-DnaA to the *in vitro* reaction resulted in a significant increase (p = 0.0004, Brown-Forsythe and Welch ANOVA tests) in the production of the reporter GFP (**Fig. 6**). Control reactions with added GST alone did not significantly change GFP levels.

**Figure 6.**
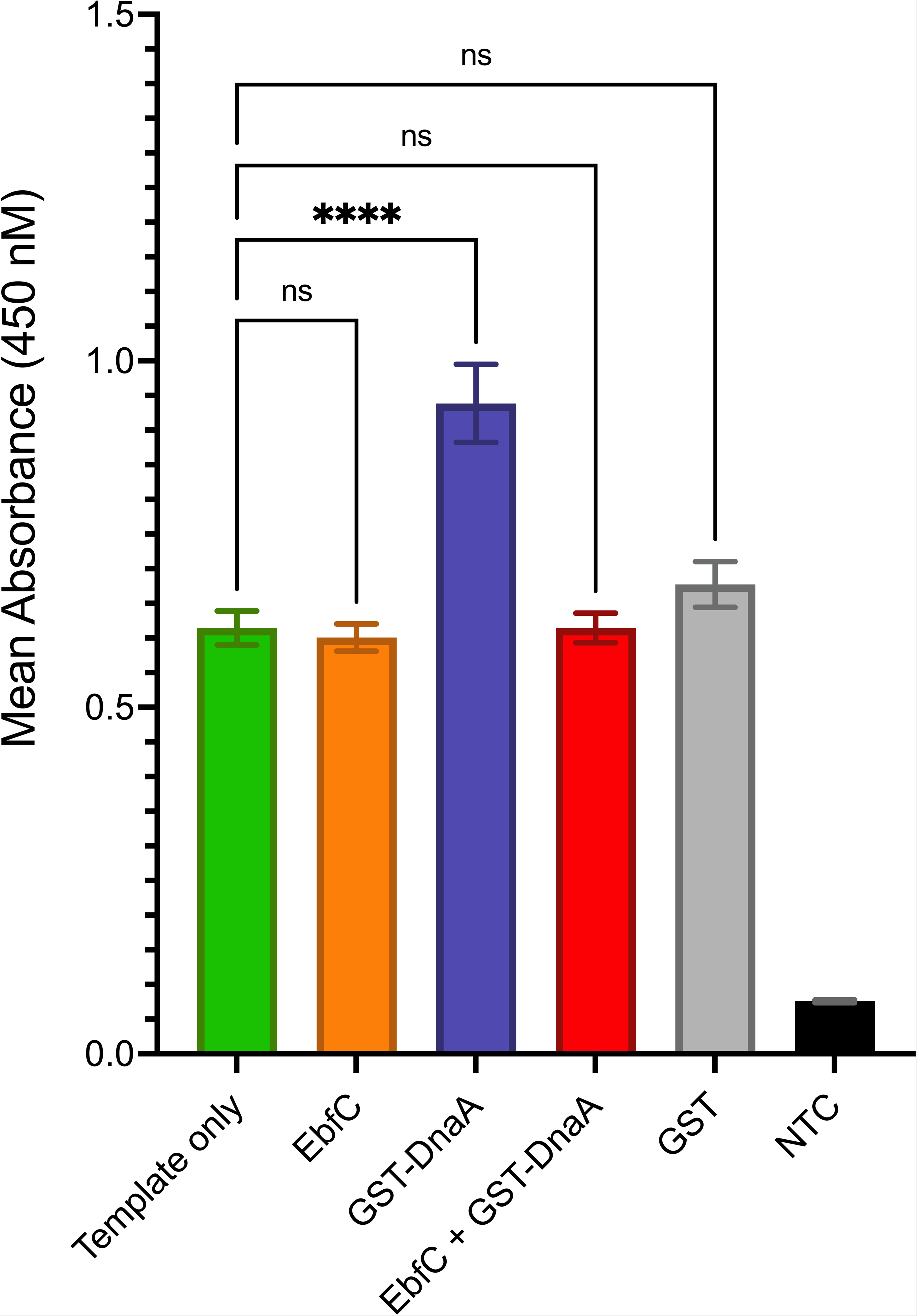
The effects of purified recombinant protein on *dnaX-ebfC in vitro* expression. Reactions used ∼65 nM linear DNA containing 247 bp upstream of the *dnaX* start codon fused to *gfp*. GST-DnaA, EbfC, or GST were added at final concentrations of 1 µM. Reporter GFP levels were quantified by ELISA and are reported as mean absorbances from triplicate reactions. The error bars indicate the standard error of the mean (SEM). Asterisks (****) indicate a statistically significant (p < 0.001) difference relative to the template-only reaction. GST served as a control to demonstrate the specific activity of the recombinant proteins. NTC = no template control.

The *in vitro* reactions with recombinant EbfC protein alone did not result in a significant change in GFP output (**Fig. 6**). Only in the presence of GST-DnaA did EbfC exhibit an effect, reducing reporter output to template-only levels (**Fig. 6**). These results indicate that *in vitro*, EbfC alone does not exert a regulatory effect, but it does inhibit the DnaA-dependent activation of *dnaX-ebfC* expression. This effect on coupled transcription and translation, combined with the apparent lack of competition between EbfC and DnaA for binding to the 5’ UTR DNA (**Fig. 5**), suggests that the effect of EbfC may be through its interaction with RNA.

### DnaA binds to 5’ DNA of other essential replication genes

The canonical bacterial DNA Pol III HE of *E. coli* consists of nine subunits. Comparatively, the borrelial replicase, like other spirochetes, has a pared-down repertoire (**Table 1**). *B. burgdorferi* has the bare minimum required to replicate its chromosome: a polymerase (DnaE, ⍺), processivity clamp (DnaN, β), clamp loader (DnaX, τ), and clamp opener (HolA, δ). *B. burgdorferi dnaX* does not appear to have the *cis*-element for translational frameshift that produces the Gamma subunit, and so might only produce the Tau subunit (*see* Supplemental material). Moreover, genes for the subunits involved with fidelity (DnaQ, ε; HolE, θ) are absent from the borrelial genome.

**Table 1.** Subunits of DNA Polymerase III Holoenzyme

| Subunit | Function | *E. coli* | *B. burgdorferi* | *T. pallidum* | *L. interrogans* |
| --- | --- | --- | --- | --- | --- |
| DnaE, $\alpha$ | Polymerase activity | B0184 | BB0579 | TP0669 | LIC10222 |
| DnaN, $\beta$ | DNA clamp; locks Pol III to DNA to increase processivity | B3701 | BB0438 | TP0002 | LIC10002 |
| DnaQ, $\epsilon$ | 3' → 5' exonuclease activity | B0215 | – | TP0643 | – |
| DnaX, $\tau$ and $\gamma$ | Dimerize core enzymes ($\tau$); clamp loader ($\gamma$) | B0470 | BB0461 | TP1005 | LIC13474 |
| HolA, $\delta$ | Clamp opener | B0640 | BB0455 | TP0588 | – |
| HolB, $\delta'$ | Clamp loader | B1099 | – | – | LIC13497 |
| HolC, $\chi$ | Interact with SSB | B4259 | – | – | – |
| HolD, $\Psi$ | Stabilize $\gamma$ and $\chi$ interaction | B4372 | – | – | – |
| HolE, $\theta$ | Proof-reading activity | B1842 | – | – | – |

Dashes (–) indicate the organism lacks an identifiable homolog.

Since we found that DnaA regulates *dnaX-ebfC*, we posited that DnaA might also be involved in regulating the other subunits of the replicase and itself. With this aim in mind, the 5’ intergenic regions of *dnaN*, *dnaE*, *holA,* and *dnaA* were used as labeled DNA probes for EMSA analysis.

The *dnaN* gene (BB_0438) codes for the processivity clamp (β) of the DNA Pol III HE and is located downstream of the *dnaA* gene (BB_0437) (**Fig. 7A**). The *B. burgdorferi oriC* is between *dnaA* and *dnaN* and contains two hypothesized DnaA-boxes (31). EMSAs demonstrated the binding of recombinant DnaA to a labeled probe that consisted of the 240 bp *dnaA-dnaN*/*oriC* region (**Fig. 7B**). The free probe was almost entirely bound by the higher levels of DnaA. There were no appreciable differences in ATP- or ADP-DnaA binding to the *oriC* probe. These results are consistent with the predicted functions of DnaA at *oriC* to control replication initiation. Further, the placement of the *oriC* in proximity to the *dnaN* promoter suggests that DnaA may regulate this gene, as it does in the firmicute *Bacillus subtilis* (32).

**Figure 7.**
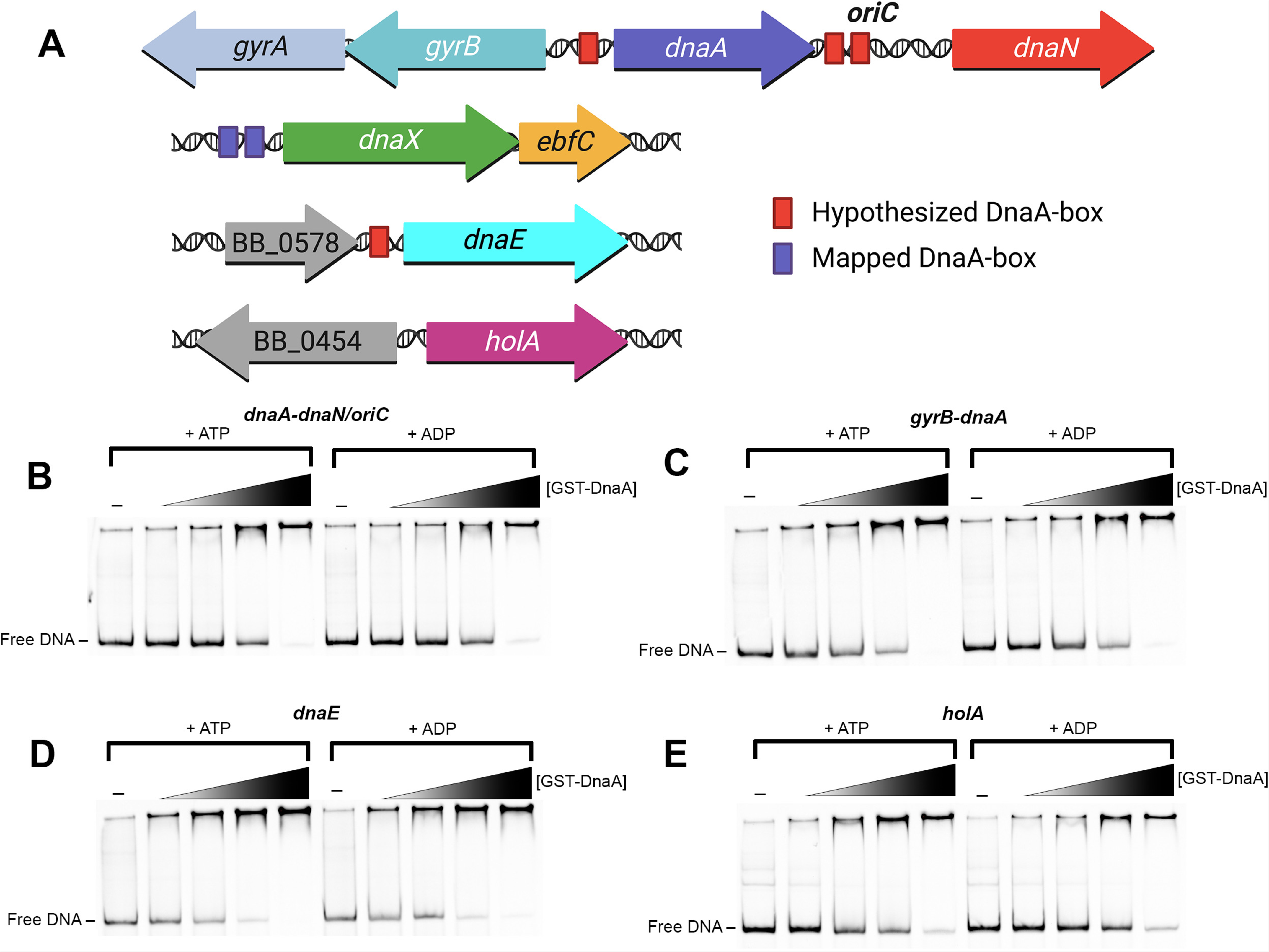
EMSAs with PCR amplified probes containing the intergenic regions 5’ of *dnaA* and the DNA pol III HE genes. (A) Schematic of the loci subject to EMSA analysis. Predicted DnaA-boxes are indicated by red rectangles. Mapped DnaA-boxes are indicated by purple rectangles. Each EMSA reaction (B-E) includes 100 nM probe, 2.5 ng/µl poly dI-dC, and either 0.56, 1.125, 2.25, and 4.5 µM GST-DnaA. The first lane for each EMSA is probe only. The first five lanes of each panel included 1 mM ATP, and the last five included 1 mM ADP. (B) Probe containing the *oriC/dnaA-dnaN* intergenic region. (C) Probe containing the intergenic region of *gyrB-dnaA*. (D) Probe containing the intergenic region 5’ of *dnaE*. (E) Probe containing the intergenic region 5’ of *holA*.

We also found that DnaA bound the 186 bp 5’ intergenic region of *dnaA* and the opposite-facing *gyrB* (BB_0436) (**Fig. 7C**), a location that contains another hypothetical DnaA-box (**Fig. 7A**) (31). The *gyrB* gene is located upstream of *gyrA* and presumably forms an operon (*gyrBA*) as in other prokaryotes. The *gyrBA* genes code for subunits A and B of the DNA gyrase, a topoisomerase essential in bacterial replication. For the *gyrB-dnaA* region, the free probe was almost entirely bound at the highest protein concentrations. ATP or ADP did not have an appreciable impact on binding to this DNA. These results show that DnaA strongly binds to its promoter region, suggesting it may undergo autoregulation as in other bacteria (25, 32–36).

The gene for the DNA polymerase DnaE (⍺, BB_0579) is located downstream of the gene for a methyl-accepting chemotaxis protein (*mcp-1*, BB_0578) (**Fig. 7A**). The intergenic region between these genes is 130 bp, and it contains a potential DnaA-box (24). EMSAs with recombinant DnaA showed binding to the *dnaE* probe (**Fig. 7D**). ATP/ADP again did not demonstrably affect binding.

The *holA* gene (δ, BB_0455) promoter overlaps with a neighboring upstream gene (BB_0454) that codes for a predicted lipid galactosyltransferase (**Fig. 7A**). This intergenic region is small, consisting of only 55 bp, and does not contain a sequence that resembles any known DnaA-box. Despite this, EMSAs showed that DnaA bound to the 5’ *holA* DNA (**Fig. 7E**).

To compare the relative affinities of DnaA for all these regions, a competition EMSA was performed using the labeled *oriC* DNA as the probe. This DNA was selected as it is the primary site for DnaA interaction and is thus an ideal standard. At 10-times molar excess, the unlabeled DNA competitors from the *dnaX-ebfC* 5’ UTR, *dnaE*, and *gyrB*-*dnaA* considerably depleted the DnaA-*oriC* complex (**Fig. 8**, lanes 5, 6, and 8, respectively). The competitor DNA from *holA* partially diminished the observed shift (**Fig. 8**, lane 7), consistent with the data above (**Fig. 7E**). The negative control DNA derived from the empty pCR2.1 vector, used for all DNAs, failed to compete (**Fig. 8**, lane 9). The unlabeled *oriC* DNA successfully depleted the DnaA-*oriC* probe complex (**Fig 8**, lane 10).

**Figure 8.**
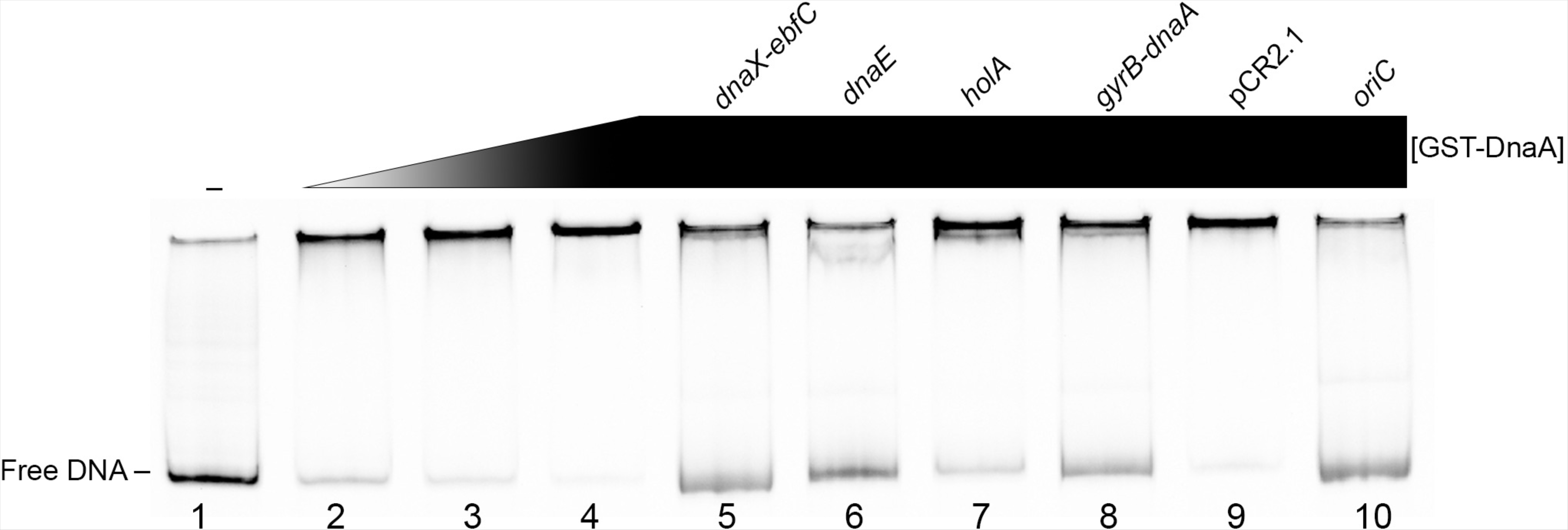
Relative affinities of DnaA for labeled *oriC* probe in comparison with other loci. Competitive EMSA using a final labeled *oriC* probe concentration of 100 nM for each lane. Lanes 2-4 have increasing concentrations of 1.125, 2.25, and 4.5 µM GST-DnaA, and lanes 5-10 have constant levels of 4.5 µM GST-DnaA. Competitor DNAs were amplified using unlabeled M13 Forward and Reverse primers from TA constructs, and added at 10-times molar excess to reactions relative to the *oriC* probe (lanes 6-10).

With knowledge of the physical interactions of DnaA at these loci, we next sought to track the transcriptional dynamics of these replication genes during the different growth phases of cultured *B. burgdorferi*. As we have established DnaA as an activator of *dnaX-ebfC* transcription *in vitro*, we hypothesized that they would have similar expression profiles. Indeed, the data show that the relative levels of *dnaX* and *ebfC* transcripts rose with *dnaA* as the culture entered the exponential phase (**Fig. 9A-C**). Moreover, *dnaX* and *dnaA* declined as the spirochetes entered the stationary growth phase, while *ebfC* remained stable for some time but eventually decreased. This divergence is consistent with our previous finding that *ebfC* has another promoter within the *dnaX* open reading frame (20).

**Figure 9.**
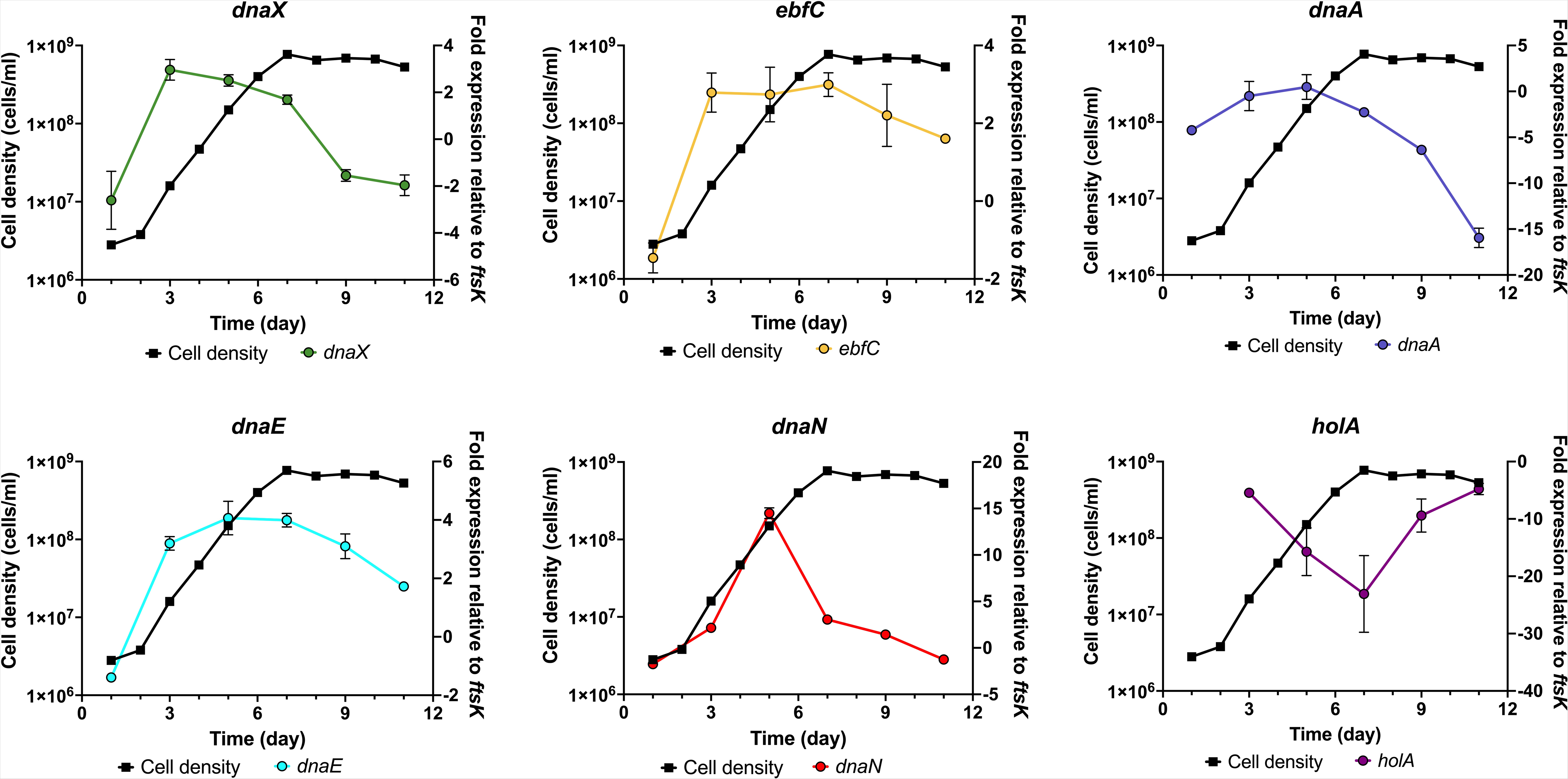
Tracking DNA Pol III HE gene expression changes during growth phases. **(A-F)** The cell density of *B. burgdorferi* (left *y*-axes) was measured every 24 h, and the relative transcript levels (right *y*-axes) were analyzed every 48 h. The fold expression of transcript was calculated by normalization to the constitutive *ftsK* gene and reported as the average of triplicate reactions. The error bars indicate the SEM.

The transcript levels of *dnaN* and *dnaE* also followed the pattern seen with *dnaA* (**Fig. 9D-E**). The relative expression level for both genes peaked at mid-exponential and steadily decreased into stationary phase. The *holA* gene, in contrast, exhibited a unique expression profile compared to the other components of the DNA Pol III HE. While the other genes’ transcripts decreased during stationary, *holA* transcript levels unexpectedly increased (**Fig. 9F**). Furthermore, the relative abundance of the *holA* transcript was considerably low, never reaching levels greater than that of the housekeeping gene *ftsK*. Those results suggest that at least one additional factor controls levels of *holA* transcript.

## DISCUSSION

Tight and coordinated regulation of chromosomal replication is a key biological process. The evolution of such regulatory networks was significant for unicellular organisms, which can inhabit dynamic environments that challenge cellular homeostasis. *B. burgdorferi* has evolved the need to concomitantly modulate DNA replication and gene expression during its consistent cycling between tick vectors and vertebrate hosts. These requirements can be appreciated at the *dnaX-ebfC* locus, where DnaX (Tau) assembles with the DNA Pol III HE to facilitate replication, while EbfC binds at sites throughout the nucleoid and presumably affects chromatin structure. Previously we reported that the expression of *dnaX-ebfC* depends on the replication rate (20, 21), but the question of how remained unanswered. This work indicates that this connection can be explained by direct interactions of DnaA and EbfC at the 5’ UTR region of *dnaX-ebfC*.

DnaA is well established to be a transcription factor in other bacterial species (19). For *B. burgdorferi*, our data show that DnaA binds 5’ of the *dnaX-ebfC* operon to enhance gene expression. This is a logical pathway to evolve, given that DnaA and DnaX are directly linked to chromosomal replication. Moreover, this mechanism is perhaps conserved as two DnaA boxes are located upstream of the *E. coli dnaX-ebfC* operon (37).

Transcriptional control of bacterial DNA replication machinery through DnaA has also been observed in *Caulobacter crescentus*. A gram-negative aquatic bacterium, *C. crescentus*, exhibits a complex cell cycle defined by asymmetric division into stalked and swarmer progeny where DNA replication is initiated only once per cell cycle (38). In *C. crescentus*, DnaA controls the transcription of two DNA replication genes, the helicase, *dnaB*, and DNA Pol III HE ε-subunit, *dnaQ* (39). There is also evidence of an unknown repressor binding to a conserved promoter motif upstream of *C. crescentus dnaX*, *dnaN*, *dnaA*, *dnaK*, and *gyrB* genes (40). Our work suggests that a similar regulation network may exist in *B. burgdorferi*, where DnaA binds upstream of *dnaX*, *dnaN*, *dnaE*, *holA*, *dnaA*, and *gyrB* (**Fig. 10**). The effect of DnaA on these borrelial genes, aside from *dnaX*, remains to be assessed.

**Figure 10.**
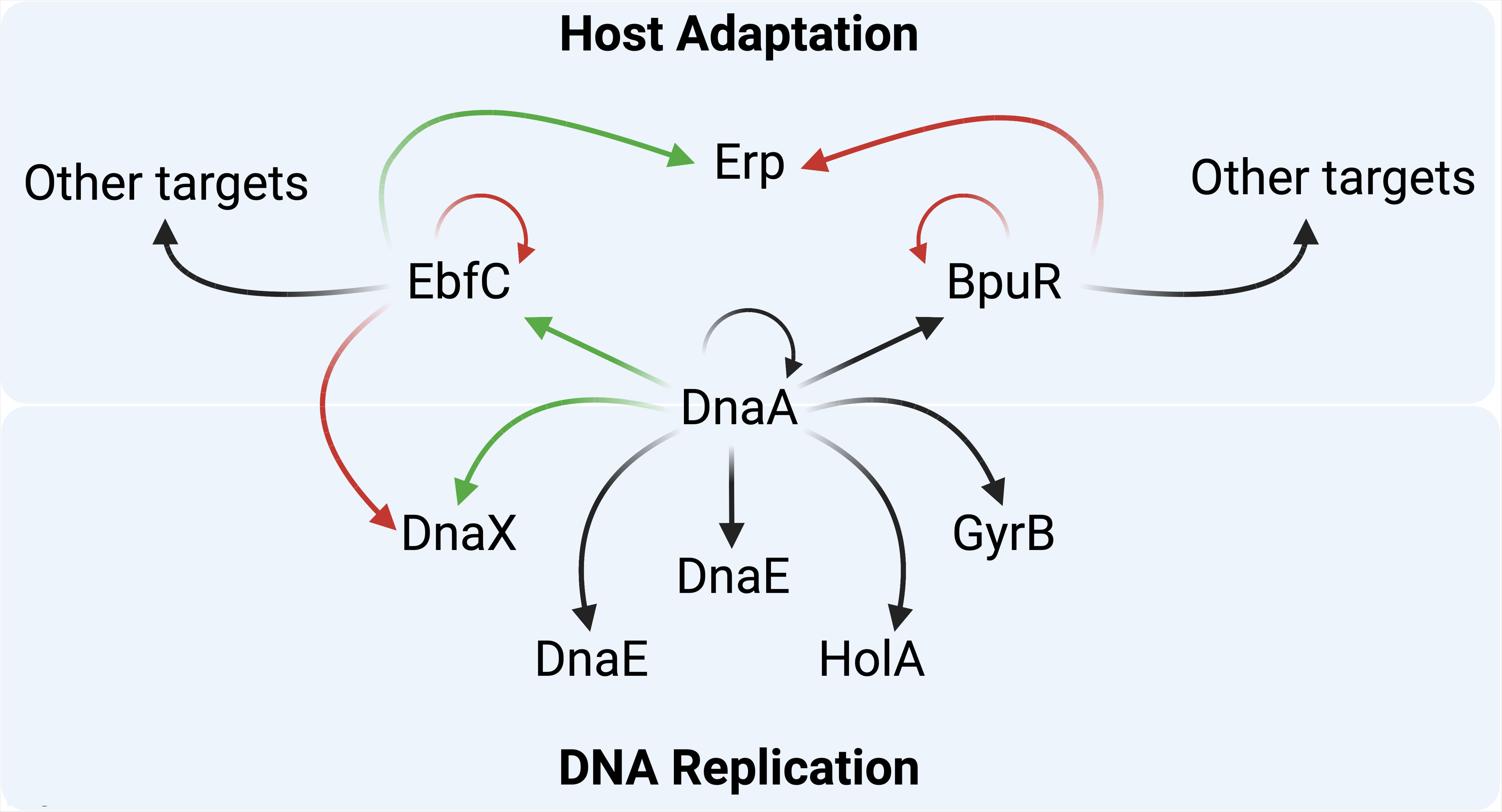
Diagram of experimentally-confirmed and hypothesized effects of DnaA on *B. burgdorferi* protein expression. Green arrows indicate positive effects, red arrows indicate negative effects, and black arrows indicate predicted effects. DnaA positively influences expression of both DnaX and EbfC. EbfC counteracts the effect of DnaA on the *dnaX-ebfC* operon, acts as an anti-repressor to stimulate transcription of *erp* operons, and also affects expression of numerous other transcripts. DnaA binds over the *bpuR* promoter and is hypothesized to repress transcription of that gene, while BpuR has been demonstrated to repress its own translation, enhance repression of *erp* operons, and affect expression of several other borrelial proteins. DnaA also binds to its own promoter region, as well as those of *dnaE, dnaN, gyrB,* and *holA*, and is hypothesized to affect expression of those operons.

The potential global transcriptional effects of DnaA in *B. burgdorferi* begs the question of what controls the master initiator. In other bacteria, the cellular ATP/ADP ratio affects the function of DnaA (25–27). Our data failed to identify a measurable effect of either ATP or ADP on DnaA’s binding to any tested sequence. This suggests either *B. burgdorferi* DnaA lacks sensitivity to those nucleotides under the tested conditions or the protein has novel control mechanisms. The latter possibility is especially intriguing due to the lack of protein homologs such as DiaA, Hda, and SeqA that regulate DnaA function/expression in other bacterial species (41–43).

The idea of novel DnaA-DNA interactions in *B. burgdorferi* has previously been hypothesized, since the borrelial homolog differs in three of eight amino acid residues in the DNA-binding C-terminus that are conserved in other species (44). In the *E. coli* DnaA, two of the divergent amino acids, T451 and S453, form base-specific hydrogen bonds with DNA (45). This suggests that the DnaA protein of *B. burgdorferi* may recognize unique DNA sequences. Our data showing DnaA binding 5’ of *holA* and *dnaX-ebfC* 5’ UTR region B, which lack sequence similarity to the characterized DnaA-boxes of other species, points to this possibility. Genome-wide distribution of bound DnaA in *B. burgdorferi* to identify the borrelial DnaA-box sequence is currently under investigation.

In addition to the regulatory activity of DnaA, our studies demonstrated that EbfC affects *dnaX-ebfC* expression. That is consistent with previous studies by our group, which found that overexpression of EbfC resulted in a decrease of *dnaX* transcript (20). The present work demonstrates that EbfC regulates *dnaX-ebfC* expression by counteracting DnaA-dependent transcriptional activation. The lack of competition between DnaA and EbfC in binding to the *dnaX-ebfC* 5’ UTR DNA suggests that EbfC may counteract the effect of DnaA by binding to *dnaX-ebfC* mRNA, which could alter secondary structure and affect transcription or translation. Further investigations to tease apart the mechanism of regulation is complicated by the fact that mutagenesis of the *dnaX-ebfC* 5’ UTR DNA will affect the structure of the mRNA.

The fluctuations from tick and vertebrate hosts have been a constant for *B. burgdorferi* for thousands of years. These divergent biological environments have forced the spirochete to evolve a regulatory network to adapt and endure appropriately. In this study, we provided evidence that *B. burgdorferi* uses its DnaA protein to promote the expression of an operon crucial to its replication and survival. DnaX, the central organizer of the DNA Pol III HE, is essential for chromosomal replication. EbfC affects expression of *dnaX-ebfC*, the vertebrate-stage outer surface Erp proteins, and many other borrelial genes. DnaA activity at *dnaX-ebfC* thus permits coordinated expression of replication machinery and a protein needed for borrelial host infection (**Fig. 10**). *B. burgdorferi*. DnaA also binds over the promoter of *bpuR*, an RNA- and DNA-binding protein that, in turn, affects expression levels of Erp and several other borrelial proteins (24). We hypothesize that *B. burgdorferi* DnaA also regulates the production of other proteins involved in replication and infection processes.

## METHODS AND MATERIALS

### Bacterial strains and culture conditions

All studies utilized derivatives of the *B. burgdorferi* type strain, B31, and were cultivated in liquid Barbour-Stoenner-Kelly II (BSK-II) medium (46). For RNA analyses, cultured bacteria were subject to temperature shift wherein spirochetes were grown at 23 °C to mid-exponential phase (∼ 5 x 10^7^ cells/ml) and passaged 1:100 into fresh BSK-II and incubated at 34 °C. Stationary phase bacteria were collected from cultures at or above 10^8^ cells/ml. Culture cell densities were determined by direct counting in a Petroff-Hauser counting chamber with darkfield microscopy.

### Recombinant proteins

EbfC was cloned into pET200 and expressed with an N-terminal 6xHis affinity tag. Recombinant EbfC was expressed in *Escherichia coli* Rosetta II cells and purified using MagneHis Particles (Promega) as previously described (24, 47).

DnaA was cloned into pET-21a(+) with an N-terminal glutathione S-transferase (GST) moiety by the University of Kentucky Protein Core. A GST tag was selected to enhance the solubility of DnaA, which in other bacteria is known to aggregate into insoluble inclusion bodies (48, 49). Recombinant GST-DnaA protein was expressed in *E. coli* BL21 DE3 cells with 1 mM isopropyl-β-d-thiogalactopyranoside (IPTG) overnight at room temperature. Recombinant GST was expressed in *E. coli* Rosetta II cells containing pGEX-3x. Cells were collected by centrifugation and frozen at −80 °C.

Frozen GST-DnaA expressing cells were resuspended in ice-cold wash/binding buffer (4.2 mM Na_2_HPO_4_, 2 mM KH_2_PO_4_, 140 mM NaCl, 10 mM KCl) and then supplemented with 5% (v/v) B-PER Bacterial Protein Extraction Reagent (Thermo Scientific) before sonication. The soluble fraction was separated by centrifugation and discarded. The insoluble fraction was retained and processed using mild solubilization per Qi and colleagues (50). Briefly, the insoluble pellet was washed three times with inclusion body wash buffer (20 mM Tris, 300 mM NaCl, 1 mM EDTA, 1% Triton X-100, 1 M urea, pH 8.0) followed by two washes with phosphate-buffered saline (PBS), pH 7.4 to remove detergent. The final pellet was resuspended in PBS with 2 M urea and frozen at −20 °C for at least 12 h. The frozen suspension was then thawed at room temperature and dialyzed at 4 °C for at least 3 h in 1 L of PBS with 1 M urea, 0.5 M urea, and no urea. The clarified soluble protein suspension was purified using MagneGST Glutathione Particles (Promega) according to manufacturer recommendations. Recombinant GST was purified using the same MagneGST conditions but under standard soluble conditions.

Purified proteins were dialyzed against EMSA binding buffer (50 mM Tris-HCl, 50 mM KCl, 1 mM EDTA, 1 mM phenylmethanesulfonyl fluoride [PMSF], 1 mM dithiothreitol [DTT], 0.01% Tween-20, 10% glycerol) overnight at 4 °C and concentrated with 10 kDa MWCO Amicon centrifugal filters (Sigma). Protein concentrations were determined by Bradford assay, and purity was assessed by SDS-PAGE with Coomassie brilliant blue staining. Final protein preparations were stored at −80 °C in 12 µl working aliquots.

### TOPO Plasmid Constructs

For these studies, DNA regions of interest were PCR amplified from *B. burgdorferi* B31 genomic DNA with Taq polymerase and TA cloned into pCR2.1 (Thermo Fisher). Constructs were sequenced to guarantee sequence fidelity.

### Electrophoretic mobility shift assay (EMSA)

Labeled and unlabeled nucleic acids used are described in the **Supplemental Table**. Oligonucleotides for these assays were synthesized by Integrated DNA Technologies (IDT, Coralville, IA). For use as EMSA probes, DNAs were conjugated with a 5’ IRDye800 fluorescent tag, while RNAs had a 5’ Alexa Fluor 647 tag. Short DNA probes (< 60 bp) were produced by heating two complementary oligonucleotides to 95 °C for 5 minutes and gradually cooling them to room temperature. Long DNAs (> 60 bp) were produced by PCR amplification from TA cloned plasmids using M13 Forward and M13 Reverse primers, with a fluor tag on the 5’ end of the M13 Reverse. PCR products were treated with exonuclease I (New England BioLabs) to remove residual primers. The reactions were precipitated by adding a 1/10 volume of 3 M sodium acetate and a 2.5 times volume of ice-cold 100% ethanol. Glycogen was added at a final concentration of 1 mg/ml to aid pellet visualization. DNA was pelleted by centrifugation at 4 °C and washed twice with ice-cold 70% ethanol to remove excess salt. The final DNA pellet was air-dried at room temperature for 15 minutes and resuspended in 50 µl of TE buffer. Nucleic acid concentrations were determined spectrophotometrically and diluted to 1 µM. Final DNA probes were stored at −20 °C in working doses of 12 µl.

EMSAs were performed essentially as described previously (11, 51). Briefly, protein-nucleic acid binding reactions were carried out at room temperature for 15 min in EMSA binding buffer using either 100 nM or 20 nM probe. All reactions with GST-DnaA were supplemented with 1 mM MgCl_2_ and 1 mM of either ATP or ADP and allowed to incubate on ice for 15 min before adding DNA. When appropriate, the non-specific competitor Poly dI dC (Roche) was added to each reaction before the probe at a final concentration of 2.5 ng/µl (52). Competitor nucleic acids were also added to reactions before the labeled probe. Prior to electrophoresis, 6x EMSA loading dye (0.8 mg/ml Orange G, 15 mg/ml Ficol 400) was added to each reaction. Electrophoresis was performed with 6% or 10% Novex TBE gels (Thermo Fisher) or in-house 6% polyacrylamide TBE gels. The in-house polyacrylamide TBE gels (40 ml) for horizontal electrophoresis were produced by combining 6 ml 40% Acryl/Bis 29:1 (VWR), 4 ml of 5x TBE, pH 7.5 (445 mM Tris, 445 mM boric acid, 5 mM EDTA), 30 ml ddH_2_O, 300 µl 10% APS (Fisher), and 100 µl of TEMED (National Diagnostic). The gels were pre-run in 0.5x TBE buffer at 12.5 V/cm for 30 min. Samples were loaded and resolved at 12.5 V/cm until the dye front reached the bottom of the gel.

The apparent K_D_ of GST-DnaA for the *dnaX-ebfC* 5’ UTR DNA probe was determined by densitometry from triplicate EMSAs using ImageLab software (BioRad). Lanes were manually set to the EMSA gel. The free DNA and shifted bands were manually selected, and the intensities were reported as lane %. These values were subsequently plotted against the corresponding protein concentration. The plotted data was then subjected to nonlinear regression analysis using Prism 9 software. An unpaired t-test was performed to assess statistically significant differences (⍺ = 0.5) between the ATP and ADP data.

### Coupled *in vitro* transcription/translation assays

Coupled *in vitro* transcription/translation assays were performed as previously described (10, 11, 30). Linear DNA template was amplified from pAAB206 using M13 forward and reverse primers (20). This amplicon contains 247 bp 5’ of *dnaX-ebfC* fused to *gfp*. Experiments were performed using the *E. coli* S30 Extract System for Linear Templates (Promega). The 50-µl reactions were composed of kit components, 2,100 ng of template DNA (∼65 nM), 40 U of RiboGuard RNase Inhibitor (Lucigen), and 1 µM of the proteins of interest, when appropriate. Purified GST was used as a control. Reactions were incubated at 37 °C for 2 h and promptly iced for 5 minutes to stop transcription/translation. Each reaction set was performed in triplicate.

The *in vitro* protein products were precipitated by adding 4-volumes of ice-cold acetone and incubating at −20 °C for at least one hour. Proteins were pelleted by centrifugation and air-dried on ice for 15 minutes. Dried protein pellets were then resuspended in 100 µl of PBS. Protein suspensions were mixed with 500 µl of ELISA coating buffer (50 mM Na_2_CO_3_, 500 mM NaHCO_3_, pH 9.2), and 150 µl of this mixture was added to the wells of 96-well microtiter plates. GFP was detected with MACS anti-GFP-HRP (Militenyi Biotec). Seventy-five microliters of tetramethyl benzidine (TMB; Thermo Scientific) HRP substrate were added to each well and incubated at room temperature with shaking for 30 minutes. Reactions were quenched with 75 µl of 2N H_2_SO_4,_ and the absorbance at 450 nm was measured using a Biotek Epoch 2 microplate reader.

Statistical significance (⍺ = 0.05) was determined using Brown-Forsythe and Welch ANOVA tests and Dunnett’s T3 multiple comparisons.

### Quantitative reverse transcription-PCR (qRT-PCR)

To measure the levels of target transcripts, *B. burgdorferi* was cultured at 35 °C, and 2 ml aliquots of bacteria were taken 1-, 3-, 5-, 7-, 9-, and 11-days post-inoculation. Cells were washed two times with PBS and frozen at −80 °C. Frozen pellets were later resuspended in 60 °C TRIzol (Thermo Fisher), and RNA was extracted using a Qiagen Mini RNA kit. Residual genomic DNA was depleted using on-column Turbo DNase (Thermo Fisher). The quality (RIN > 7) and quantity of the RNA were determined using an Agilent 2100 Bioanalyzer and Agilent 6000 Nano chips. Purified RNA stocks were stored at −80 °C.

Complementary DNA (cDNA) for qPCR was produced using the iScript gDNA Clear cDNA synthesis kit (Bio-Rad). The cDNA stocks were diluted 1:100 into nuclease-free water for use as template. Quantitative PCR was carried out using iTaq Universal SYBR Green Supermix (Bio-Rad) and a Bio-Rad CFX96 Touch Real-time PCR thermocycler. Oligonucleotide primer sets for target genes were designed using the IDT Primer Quest Tool (https://www.idtdna.com/PrimerQuest/Home/Index). For all reactions, cycles involved melting at 95 °C for 10s, annealing at 55 °C for 10 s, and extension at 72 °C for 30s. Each run was performed with technical triplicates of each reaction as well as no template control reactions that lacked template. Melting curves were performed with each run to validate the production of specific amplicons. Every reaction was tested with a reverse transcriptase-negative template to validate the absence of contaminating gDNA. Cq values were normalized to *ftsK*, a gene whose expression is stable across different growth phases (53), using the ΔCq method, and fold-difference was determined by the function 2^−Δ^ ^Cq^.

## Acknowledgments

This research was funded by NIH grant R21 AI147139-02. We thank Tatiana Castro-Padovani and Nerina Jusufovic for their helpful comments and support on these studies and the manuscript. The construct for recombinant GST-DnaA was synthesized by Drs. Martin Chow and Lou Hersh of the University of Kentucky COBRE Protein Core. Dr. Tony Sinai kindly provided pGEX-3x. Schematic figures were made using BioRender.

## Works Cited

1. Kawakami H, Katayama T. 2010. DnaA, ORC, and Cdc6: similarity beyond the domains of life and diversity. Biochem Cell Biol 88:49–62.

2. McHenry CS. 2011. DNA replicases from a bacterial perspective. Annu Rev Biochem 80:403–36.

3. Speck C, Messer W. 2001. Mechanism of origin unwinding: sequential binding of DnaA to double- and single-stranded DNA. EMBO J 20:1469–76.

4. Kaur G, Vora MP, Czerwonka CA, Rozgaja TA, Grimwade JE, Leonard AC. 2014. Building the bacterial orisome: high-affinity DnaA recognition plays a role in setting the conformation of *oriC* DNA. Mol Microbiol 91:1148–63.

5. Ozaki S, Katayama T. 2012. Highly organized DnaA-*oriC* complexes recruit the single-stranded DNA for replication initiation. Nucleic Acids Res 40:1648–65.

6. Dunham-Ems SM, Caimano MJ, Pal U, Wolgemuth CW, Eggers CH, Balic A, Radolf JD. 2009. Live imaging reveals a biphasic mode of dissemination of *Borrelia burgdorferi* within ticks. J Clin Invest 119:3652–65.

7. Piesman J, Oliver JR, Sinsky RJ. 1990. Growth kinetics of the Lyme disease spirochete (*Borrelia burgdorferi*) in vector ticks (*Ixodes dammini*). Am J Trop Med Hyg 42:352–7.

8. De Silva AM, Fikrig E. 1995. Growth and migration of *Borrelia burgdorferi* in *Ixodes* ticks during blood feeding. Am J Trop Med Hyg 53:397–404.

9. Stevenson B, Seshu J. 2018. Regulation of Gene and Protein Expression in the Lyme Disease Spirochete. Curr Top Microbiol Immunol 415:83–112.

10. Jutras BL, Verma A, Adams CA, Brissette CA, Burns LH, Whetstine CR, Bowman A, Chenail AM, Zuckert WR, Stevenson B. 2012. BpaB and EbfC DNA-binding proteins regulate production of the Lyme disease spirochete’s infection-associated Erp surface proteins. J Bacteriol 194:778–86.

11. Jutras BL, Chenail AM, Carroll DW, Miller MC, Zhu H, Bowman A, Stevenson B. 2013. Bpur, the Lyme disease spirochete’s PUR domain protein: identification as a transcriptional modulator and characterization of nucleic acid interactions. J Biol Chem 288:26220–34.

12. Blinkova A, Hervas C, Stukenberg PT, Onrust R, O’Donnell ME, Walker JR. 1993. The *Escherichia coli* DNA polymerase III holoenzyme contains both products of the *dnaX* gene, tau and gamma, but only tau is essential. J Bacteriol 175:6018–27.

13. Dallmann HG, Thimmig RL, McHenry CS. 1995. DnaX complex of *Escherichia coli* DNA polymerase III holoenzyme. Central role of tau in initiation complex assembly and in determining the functional asymmetry of holoenzyme. J Biol Chem 270:29555–62.

14. Cordeiro TFVB, Gontijo MTP, Jorge GP, Brocchi M. 2022. EbfC/YbaB: A Widely Distributed Nucleoid-Associated Protein in Prokaryotes. Microorganisms 10:1945.

15. Tsuchihashi Z. 1991. Translational frameshifting in the *Escherichia coli dnaX* gene *in vitro*. Nucleic Acids Res 19:2457–62.

16. Tsuchihashi Z, Brown PO. 1992. Sequence requirements for efficient translational frameshifting in the *Escherichia coli dnaX* gene and the role of an unstable interaction between tRNA(Lys) and an AAG lysine codon. Genes Dev 6:511–9.

17. Larsen B, Gesteland RF, Atkins JF. 1997. Structural probing and mutagenic analysis of the stem-loop required for *Escherichia coli dnaX* ribosomal frameshifting: programmed efficiency of 50%. J Mol Biol 271:47–60.

18. Messer W, Weigel C. 1997. DnaA initiator--also a transcription factor. Mol Microbiol 24:1–6.

19. Menikpurage IP, Woo K, Mera PE. 2021. Transcriptional Activity of the Bacterial Replication Initiator DnaA. Front Microbiol 12:662317.

20. Jutras BL, Bowman A, Brissette CA, Adams CA, Verma A, Chenail AM, Stevenson B. 2012. EbfC (YbaB) is a new type of bacterial nucleoid-associated protein and a global regulator of gene expression in the Lyme disease spirochete. J Bacteriol 194:3395–406.

21. Jutras BL, Chenail AM, Stevenson B. 2013. Changes in bacterial growth rate govern expression of the *Borrelia burgdorferi* OspC and Erp infection-associated surface proteins. J Bacteriol 195:757–64.

22. Solovyev V, Salamov A, Li R. 2011. Metagenomics and its applications in agriculture, biomedicine and environmental studies. Autom Annot Microb Genomes Metagenomic Seq Nova Science.

23. Adams PP, Flores Avile C, Popitsch N, Bilusic I, Schroeder R, Lybecker M, Jewett MW. 2017. *In vivo* expression technology and 5’ end mapping of the *Borrelia burgdorferi* transcriptome identify novel RNAs expressed during mammalian infection. Nucleic Acids Res 45:775–792.

24. Jutras BL, Savage CR, Arnold WK, Lethbridge KG, Carroll DW, Tilly K, Bestor A, Zhu H, Seshu J, Zuckert WR, Stewart PE, Rosa PA, Brissette CA, Stevenson B. 2019. The Lyme disease spirochete’s BpuR DNA/RNA-binding protein is differentially expressed during the mammal-tick infectious cycle, which affects translation of the SodA superoxide dismutase. Mol Microbiol 112:973–991.

25. Speck C, Weigel C, Messer W. 1999. ATP- and ADP-*dnaA* protein, a molecular switch in gene regulation. EMBO J 18:6169–76.

26. Smith JL, Grossman AD. 2015. *In Vitro* Whole Genome DNA Binding Analysis of the Bacterial Replication Initiator and Transcription Factor DnaA. PLoS Genet 11:e1005258.

27. Sekimizu K, Bramhill D, Kornberg A. 1987. ATP activates *dnaA* protein in initiating replication of plasmids bearing the origin of the *E. coli* chromosome. Cell 50:259–65.

28. Riley SP, Bykowski T, Cooley AE, Burns LH, Babb K, Brissette CA, Bowman A, Rotondi M, Miller MC, DeMoll E, Lim K, Fried MG, Stevenson B. 2009. *Borrelia burgdorferi* EbfC defines a newly-identified, widespread family of bacterial DNA-binding proteins. Nucleic Acids Res 37:1973–83.

29. Van Gundy T, Lybecker M. Identification of RNA chaperones by gradient profiling in the Lyme disease spirochete.

30. Jutras BL, Jones GS, Verma A, Brown NA, Antonicello AD, Chenail AM, Stevenson B. 2013. Posttranscriptional self-regulation by the Lyme disease bacterium’s BpuR DNA/RNA-binding protein. J Bacteriol 195:4915–23.

31. Picardeau M, Lobry JR, Hinnebusch BJ. 1999. Physical mapping of an origin of bidirectional replication at the centre of the *Borrelia burgdorferi* linear chromosome. Mol Microbiol 32:437–45.

32. Ogura Y, Imai Y, Ogasawara N, Moriya S. 2001. Autoregulation of the *dnaA-dnaN* operon and effects of DnaA protein levels on replication initiation in *Bacillus subtilis*. J Bacteriol 183:3833–41.

33. Berenstein D, Olesen K, Speck C, Skovgaard O. 2002. Genetic organization of the *Vibrio harveyi* DnaA gene region and analysis of the function of the *V. harveyi* DnaA protein in *Escherichia coli*. J Bacteriol 184:2533–8.

34. Salazar L, Guerrero E, Casart Y, Turcios L, Bartoli F. 2003. Transcription analysis of the *dnaA* gene and *oriC* region of the chromosome of *Mycobacterium smegmatis* and *Mycobacterium bovis* BCG, and its regulation by the DnaA protein. Microbiology (Reading) 149:773–784.

35. Jakimowicz D, Majka J, Lis B, Konopa G, Wegrzyn G, Messer W, Schrempf H, Zakrzewska-Czerwinska J. 2000. Structure and regulation of the *dnaA* promoter region in three *Streptomyces* species. Mol Gen Genet 262:1093–102.

36. Atlung T, Clausen E, Hansen FG. 1984. Autorepression of the *dnaA* gene of *Escherichia coli*. Adv Exp Med Biol 179:199–207.

37. Flower AM, McHenry CS. 1986. The adjacent *dnaZ* and *dnaX* genes of *Escherichia coli* are contained within one continuous open reading frame. Nucleic Acids Res 14:8091–101.

38. Collier J. 2019. Cell division control in *Caulobacter crescentus*. Biochim Biophys Acta Gene Regul Mech 1862:685–690.

39. Hottes AK, Shapiro L, McAdams HH. 2005. DnaA coordinates replication initiation and cell cycle transcription in *Caulobacter crescentus*. Mol Microbiol 58:1340–53.

40. Keiler KC, Shapiro L. 2001. Conserved promoter motif is required for cell cycle timing of *dnaX* transcription in *Caulobacter*. J Bacteriol 183:4860–5.

41. Ishida T, Akimitsu N, Kashioka T, Hatano M, Kubota T, Ogata Y, Sekimizu K, Katayama T. 2004. *DiaA*, a novel DnaA-binding protein, ensures the timely initiation of *Escherichia coli* chromosome replication. J Biol Chem 279:45546–55.

42. Kato J, Katayama T. 2001. Hda, a novel DnaA-related protein, regulates the replication cycle in *Escherichia coli*. EMBO J 20:4253–62.

43. Taghbalout A, Landoulsi A, Kern R, Yamazoe M, Hiraga S, Holland B, Kohiyama M, Malki A. 2000. Competition between the replication initiator DnaA and the sequestration factor SeqA for binding to the hemimethylated chromosomal origin of *E. coli in vitro*. Genes to Cells 5:873–884.

44. Old IG, Margarita D, Saint Girons I. 1993. Unique genetic arrangement in the *dnaA* region of the *Borrelia burgdorferi* linear chromosome: nucleotide sequence of the *dnaA* gene. FEMS Microbiol Lett 111:109–14.

45. Fujikawa N, Kurumizaka H, Nureki O, Terada T, Shirouzu M, Katayama T, Yokoyama S. 2003. Structural basis of replication origin recognition by the DnaA protein. Nucleic Acids Res 31:2077–86.

46. Barbour AG. 1984. Isolation and cultivation of Lyme disease spirochetes. Yale J Biol Med 57:521–5.

47. Savage CR, Jutras BL, Bestor A, Tilly K, Rosa PA, Tourand Y, Stewart PE, Brissette CA, Stevenson B. 2018. *Borrelia burgdorferi* SpoVG DNA- and RNA-Binding Protein Modulates the Physiology of the Lyme Disease Spirochete. J Bacteriol 200.

48. Sutton MD, Kaguni JM. 1997. Threonine 435 of *Escherichia coli* DnaA protein confers sequence-specific DNA binding activity. J Biol Chem 272:23017–24.

49. Zawilak-Pawlik AM, Kois A, Zakrzewska-Czerwinska J. 2006. A simplified method for purification of recombinant soluble DnaA proteins. Protein Expr Purif 48:126–33.

50. Qi X, Sun Y, Xiong S. 2015. A single freeze-thawing cycle for highly efficient solubilization of inclusion body proteins and its refolding into bioactive form. Microb Cell Fact 14:24.

51. Babb K, Bykowski T, Riley SP, Miller MC, Demoll E, Stevenson B. 2006. Borrelia burgdorferi EbfC, a novel, chromosomally encoded protein, binds specific DNA sequences adjacent to *erp* loci on the spirochete’s resident cp32 prophages. J Bacteriol 188:4331–9.

52. Laniel MA, Béliveau A, Guérin SL. 2001. Electrophoretic mobility shift assays for the analysis of DNA-protein interactions. Methods Mol Biol 148:13–30.

53. Arnold WK, Savage CR, Brissette CA, Seshu J, Livny J, Stevenson B. 2016. RNA-Seq of *Borrelia burgdorferi* in Multiple Phases of Growth Reveals Insights into the Dynamics of Gene Expression, Transcriptome Architecture, and Noncoding RNAs. PLoS One 11:e0164165.

