## Supplementary material for "*Borrelia burgdorferi* DnaA and the nucleoid-associated protein EbfC coordinate expression of the *dnaX-ebfC* operon": S Figure (from another version of this preprint: version 1, 2022-10-18)

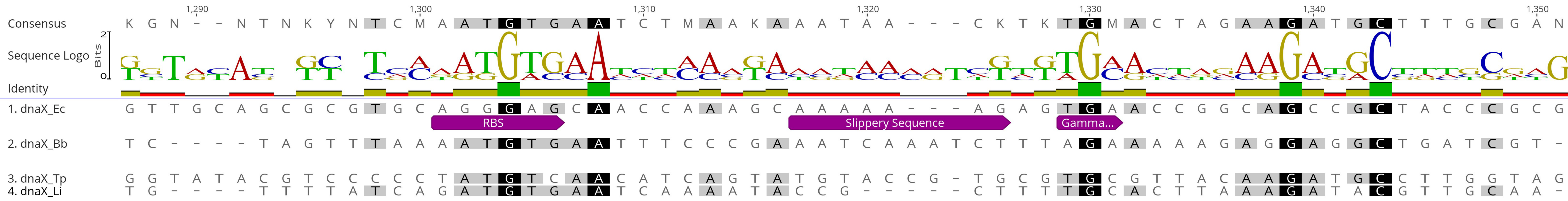


% Gene Identity

| | ***dnaX Ec*** | ***dnaX Bb*** | ***dnaX Tp*** | ***dnaX Li*** |
|---|---|---|---|---|
| ***dnaX Ec*** | – | 36.546% | 36.947% | 38.413% |
| ***dnaX Bb*** | 36.546% | – | 43.739% | 46.619% |
| ***dnaX Tp*** | 36.947% | 43.739% | – | 37.076% |
| ***dnaX Li*** | 38.413% | 46.619% | 37.076% | – |

% Protein Identity

| | **DnaX *Ec*** | **DnaX *Bb*** | **DnaX *Tp*** | **DnaX *Li*** |
|---|---|---|---|---|
| **DnaX *Ec*** | – | 22.152% | 22.946% | 25.093% |
| **DnaX *Bb*** | 22.152% | – | 31.933% | 31.213% |
| **DnaX *Tp*** | 22.946% | 31.933% | – | 26.233% |
| **DnaX *Li*** | 25.093% | 31.213% | 26.233% | – |
