## Supplementary material for "*Borrelia burgdorferi* DnaA and the nucleoid-associated protein EbfC coordinate expression of the *dnaX-ebfC* operon": S Table (from another version of this preprint: version 1, 2022-10-18)

| **Table 2. Oligonucleotides used in this study.** | | | |
| --- | --- | --- | --- |
| **Target** | **Name** | **Sequence** | **Purpose** |
| *dnaX-ebfC* 5’ UTR | *dnaX* F | GGTTTAATTTTGAACCCAA | EMSA probe 1, clone pJST01 |
|  | *dnaX* R* | CCTCCCTATGCTTTAATAAG |  |
| *dnaA* promoter | P *dnaA* F | TAGCAACATAATTCATAAACCTTCCA | Clone pACK100 |
|  | P *dnaA* R | TTTGTCCTTGTTGGTTTTAGCACT |  |
| *oriC*/*dnaN* promoter | *oriC* F | GATAGAGACAGAAATAATGACAAAG | Clone pACK101 |
|  | *oriC* R | GTTGTGTAGCATGATGCCTC |  |
| *dnaE* promoter | P *dnaE* F | ATGGTAATTGAATTGGATAATCAAGGA | Clone pACK107 |
|  | P *dnaE* R | ACCTAGACCTAAAACTCATACCTAAA |  |
| *holA* promoter | P *holA* F | GGATATATGTATCTGTAAATATTGC | Clone pACK111 |
|  | P *holA* R | CCAACAATAAATAAACCGC |  |
| pCR2.1 backbone | M13 F | CAGGAAACAGCTATGAC | PCR TA probes |
|  | M13 R* | GTAAAACGACGGCCAGT |  |
| *dnaA* ORF | qPCR *dnaA* F | CATGTGACCGATCTCCTTCTG | qPCR |
|  | qPCR *dnaA* R | CGACAATAGCTGCTCTGAGTT |  |
| *dnaX* ORF | qPCR *dnaX* F | GTGGTAGCGTAAGAGATGCTTAT | qPCR |
|  | qPCR *dnaX* R | GGTTAAGCCCATCTTGGATCT |  |
| *ebfC* ORF | qPCR *ebfC* F | TGTGGTAAAGCAGGTAGCAATA | qPCR |
|  | qPCR *ebfC* R | GAGACAGCATCATTTAAAGCAGAT |  |
| *dnaN* ORF | qPCR *dnaN* F | GTCGGGAGAAGGTATGGTTAAA | qPCR |
|  | qPCR *dnaN* R | CTCATCTGCTCCGTCATACAG |  |
| *dnaE* ORF | qPCR *dnaE* F | CATTGGTTCTTGAGGGAATGAATAG | qPCR |
|  | qPCR *dnaE* R | AGAGGCACATAGTCGGTTAAAG |  |
| *holA* ORF | qPCR *holA* F | CAGCTGTAGGATTTGCTGAGA | qPCR |
|  | qPCR *holA* R | GCTATATCTTCCTCCTCAATGGTATTA |  |
| *ftsK* ORF | qPCR *ftsK* F | GACCTTCTGATGAGCCAATGT | qPCR |
|  | qPCR *ftsk* R | GCTGCTCTGTTGTAACCTATCT |  |
| Consensus EbfC motif | EbfC F* | GGATTTGTCTTTTTGTAACTTTCAATTTTA | EMSA probe 2 |
|  | EbfC R | TAAAATTGAAAGTTACAAAAAGACAAATCC |  |
| Partial EbfC | DnaA F* | TCATAAAATTCTTTTAAAGTAATAGCTCTG | EMSA probe 3 |
|  | DnaA R | CAGAGCTATTACTTTAAAAGAATTTTATGA |  |
| *dnaX-ebfC* 5’ UTR | DPC1 F | TTAATTTTGAACCCAAATGCAATGTTTTGCTTTATGTTAAGATAT | *dnaX-ebfC* 5’ UTR Competitor 1 |
|  | DPC1 R | ATATCTTAACATAAAGCAAAACATTGCATTTGGGTTCAAAATTAA |  |
| *dnaX-ebfC* 5’ UTR | DPC2 F | GGATTTGTCTTTTTGTAACTTTCAATTTTATTATATTTTTTATGT | *dnaX-ebfC* 5’ UTR Competitor 2 |
|  | DPC2 R | ACATAAAAAATATAATAAAATTGAAAGTTACAAAAAGACAAATCC |  |
| *dnaX-ebfC* 5’ UTR | DPC3 F | TTCATAAAATTCTTTTAAAGTAATAGCTCTGAGATATAAATATGT | *dnaX-ebfC* 5’ UTR Competitor 3 |
|  | DPC3 R | ACATATTTATATCTCAGAGCTATTACTTTAAAAGAATTTTATGAA |  |
| *dnaX-ebfC* 5’ UTR | DPC4 F | TGTATAGTATATATTTACTAATATTTTACTTATTAAAGCATAGGG | *dnaX-ebfC* 5’ UTR Competitor 4 |
|  | DPC4 R | CCCTATGCTTTAATAAGTAAAATATTAGTAAATATATACTATACA |  |
| *dnaX-ebfC* 5’ UTR | DPC5 F | AAATGCAATGTTTTGCTTTATGTTAAGATATGGATTTGTCTTTTT | *dnaX-ebfC* 5’ UTR Competitor 5 |
|  | DPC5 R | AAAAAGACAAATCCATATCTTAACATAAAGCAAAACATTGCATTT |  |
| *dnaX-ebfC* 5’ UTR | DPC6 F | TTTCAATTTTATTATATTTTTTATGTTTCATAAAATTCTTTTAAA | *dnaX-ebfC* 5’ UTR Competitor 6 |
|  | DPC6 R | TTTAAAAGAATTTTATGAAACATAAAAAATATAATAAAATTGAAA |  |
| *dnaX-ebfC* 5’ UTR | DPC7 F | AGCTCTGAGATATAAATATGTTGTATAGTATATATTTACTAATAT | *dnaX-ebfC* 5’ UTR Competitor 7 |
|  | DPC7 R | ATATTAGTAAATATATACTATACAACATATTTATATCTCAGAGCT |  |
| Consensus EbfC motif | EbfC RNA** | GGAUUUGUCUUUUUGUAACUUUCAAUUUUA | EMSA RNA probe & competitor |
| *flaB* mRNA | *flaB* RNA | AGGCAAAAGGAUUUGCCAAAGUCAGAAAUU | EMSA RNA competitor |

* Oligo for probe conjugated with an IRDye800 fluorescent tag.

** Oligo for probe conjugated with an Alexa 647 fluorescent tag.
